# The technical-reasoning network is recruited when people observe others make or teach how to make tools: An fMRI study

**DOI:** 10.1101/2024.03.21.586121

**Authors:** Alexandre Bluet, Emanuelle Reynaud, Giovanni Federico, Chloé Bryche, Mathieu Lesourd, Arnaud Fournel, Franck Lamberton, Danielle Ibarrola, Yves Rossetti, François Osiurak

## Abstract

Cumulative technological culture is defined as the increase in efficiency and complexity of tools and techniques over generations. While the role of social cognitive skills in cultural transmission has been long acknowledged, recent accounts have emphasized that non-social cognitive skills such as technical reasoning, a form of causal reasoning aimed at understanding the physical world, are also at work during the social transmission of technical content. Here we contribute to this double process approach by reporting an fMRI study about the neurocognitive origins of social learning. Participants were shown videos depicting tool-making episodes in three social-learning conditions: Reverse engineering, Observation and Teaching. Our results showed that the technical-reasoning network, centred around the Area PF of the left inferior parietal cortex, was preferentially activated when watching tool-making episodes. Additionally, the teaching component was related to an activation of the right middle temporal gyrus. We propose that technical reasoning is at the heart of our technological culture and that the role of social cognition and teaching is to improve the learner’s technical reasoning by helping them concentrate on important parts of the technology. Thus, both technical reasoning and social-cognitive skills may play a key role in the cultural evolution of our technologies.

## Introduction

The increasing efficiency and complexity of tools and toolmaking techniques over generations (i.e., cumulative technological culture, hereafter shortened as CTC) enable humans to be successful ecologically and demographically, allowing them to expand all over the world, even beyond Earth (Boyd & Richerson, 1985; Derex, 2022; Tomasello et al., 1993). CTC depends on the transmission of technical knowledge, which is supported by social learning. Social learning refers to learning about conspecifics or the physical world that is influenced by observation of, or interaction with, another conspecific or its products (Heyes, 1994). Technological culture can be seen in nonhuman animals [henceforth ‘animals’; for a review, see (Whiten, 2021)], but human technological culture differs because of its cumulative component, which results in products that could not have been invented by a single individual (Tomasello, 1999) although evidence for non-technological cumulative culture as been shown in animals (Sasaki & Biro, 2017; Whiten et al., 2022). This distinction between animal and human technological culture has led researchers over the years to investigate the origins (i.e., the basis) of CTC. In the past four decades, important insights have been gained from disciplines such as evolutionary biology, mathematics, anthropology, archaeology, economics, and some psychological sub-disciplines such as social, developmental, or comparative psychology. This contrasts with the smaller contribution of cognitive sciences and particularly cognitive neuroscience (Heyes, 2018b, 2018a). Here we contribute to fill this lack by reporting an fMRI study about the neurocognitive origins of social learning during tool-making episodes.

CTC is driven by two engines, namely, high-fidelity transmission and innovation (Legare & Nielsen, 2015). The “ratcheting” hypothesis places heavy emphasis on the high-fidelity transmission component (Tennie et al., 2009; Tomasello, 1999; Tomasello et al., 1993). It assumes that cumulative culture can emerge only because high-fidelity transmission maintains cultural traits in place between innovative events. This hypothesis has led researchers for the last decades to turn their attention to the social part of CTC and to make fundamental discoveries [e.g., over-imitation (Hoehl et al., 2019; Horner & Whiten, 2005; Lyons et al., 2007), distinct forms of teaching (Kline, 2015) and social-learning strategies (Kendal et al., 2018)] about the social cognitive skills, namely mentalizing, the ability to detect as well as attribute intentions and mentals states in others (2), that can support high-fidelity transmission. Without denying the role of mentalizing, recent accounts have perceived an over-emphasis on the social dimension of the phenomenon and have argued that substantial effort must also be devoted to understanding the non-social cognitive skills at work in both the social and asocial episodes of transmission (Osiurak, Claidière, & Federico, 2022; Osiurak & Reynaud, 2020; Perry et al., 2021; Singh et al., 2021; Vale et al., 2021; Whiten, 2022).

The technical-reasoning hypothesis is consistent with these accounts (Osiurak, Claidière, & Federico, 2022; Osiurak & Reynaud, 2020). Technical reasoning is a specific form of causal reasoning oriented towards the physical world. Technical reasoning is based on mechanical knowledge that is acquired through experience (i.e., asocial and social learning). It allows humans to understand how tools are used but also how they are built, and thus increases the fidelity of transmitting such information to another peer. This hypothesis is supported by experimental work using microsociety paradigms (e.g., experiments where chains of individuals are asked to improve a physical system), which have shown that technical-reasoning skills are a key predictor of cumulative performance (De Oliveira et al., 2019; Osiurak et al., 2016, 2020) or that the improvement of a physical system over generations is accompanied by an increased understanding of it [(Osiurak, Claidière, Bluet, et al., 2022; Osiurak, Lasserre, et al., 2021) but see (Derex et al., 2019; Kendal, 2019); for a review see (Osiurak, Claidière, & Federico, 2022)]. According to this hypothesis (Bluet et al., 2022; Osiurak & Reynaud, 2020), mentalizing is primarily involved in teaching episodes, that is, when the model modifies their behavior to facilitate learning in others (Caro & Hauser, 1992; Kline, 2015). In these episodes, mentalizing acts to strengthen and improve reasoning, the teacher pointing to where the focus of reasoning should be put. Neuropsychological and neuroimaging evidence (Goldenberg & Spatt, 2009; Reynaud et al., 2016) has indicated that the human technical-reasoning network is primarily comprised of a specific region located in the left inferior parietal lobe (IPL), named area PF (Parietal F), and of the left Inferior Frontal Gyrus (IFG). This network is recruited not only when people are reasoning about tool-use (Reynaud et al., 2016) or physical events (Fischer et al., 2016), but also when they are watching others use tools (Reynaud et al., 2019), which confirms that technical reasoning could be involved in both asocial and social learning.

The goal of the present study was to investigate the neurocognitive origins of social learning during tool-making episodes. In this paper, we define a tool as an object from the environment that is used to perform a goal-directed action (Mangalam et al., 2022). This definition encompass both what Schumaker call tool and also construction (Shumaker et al., 2011). This study was motivated by two reasons. First, CTC, strictly speaking, describes the transmission of tool-making techniques (e.g., a bow) and not of tool-use techniques. Although most of the tool-making techniques involve the use of additional tools, it remains that showing that the technical-reasoning network is recruited when someone is observing others use tools does not demonstrate that this network is recruited when someone is watching others make tools. Thus, investigating the neurocognitive bases of the “tool-making observation network” may help us draw a more direct link between technical reasoning, tool making, social learning and, as a result, CTC. Second, the technical-reasoning hypothesis predicts that technical reasoning is recruited in all social-learning episodes in which technical content is transmitted, whereas mentalizing is specifically recruited in teaching episodes. Again, investigating the neurocognitive bases of the “tool-making observation network” may be useful to test this prediction.

To do so, we presented participants with videos depicting tool-making episodes (three experimental conditions) or item-transport episodes (i.e., control condition; **Fig. 1**). The participants had to watch the video and respond to whether the video depicted a tool-making episode or not. The three experimental social-learning conditions were as follows. The first was Reverse engineering, where you watch a tool and try, by yourself, to infer how it was made. In the present study, participants watched a model showing the tool at three different steps of its making. The second was Observation, where participants watched a model make a tool, but the model did not modify his behavior to facilitate learning in participants. The third was Teaching, where participants watched a model make a tool and the model modified his behavior to facilitate learning in participants (e.g., pointing out a crucial part of the tool). Our predictions were as follows. Our first prediction was that when someone watches others make tools, technical reasoning is at work. So, its network (i.e., left area PF and left IFG) should be activated in the three social-learning conditions. Our second prediction was that mentalizing should be at play only in the Teaching condition. Mentalizing skills involve a network [(Gallagher & Frith, 2003) see also (Gallagher & Frith, 2003; Molenberghs et al., 2016; Van Overwalle & Baetens, 2009)] constituted of the anterior cingulate cortex (ACC; perspective taking), the temporal pole (TP; knowledge about persons), and the superior temporal sulcus (STS; biological motion). Additionally, several studies have revealed the importance of the right middle temporal gyrus (MTG) in the detection and understanding of communicative gestures (Holler et al., 2015) as well as communicative intent (Redcay et al., 2016), which is deemed crucial for mentalizing skills (O’Madagain & Tomasello, 2022). Besides, the right MTG is part of the mentalizing network (Hodgson et al., 2022). Hence, we predicted that the right MTG should be preferentially activated in the Teaching condition. Finally, we also expected that our social-learning conditions recruited additional brain areas involved in the tool-use observation network but also more generally in the action observation network [i.e., tool-use∩action observation network; (Reynaud et al., 2019)], namely the intraparietal sulcus (IPS), the inferior temporal gyrus (ITG), and the left middle frontal gyrus (MFG).

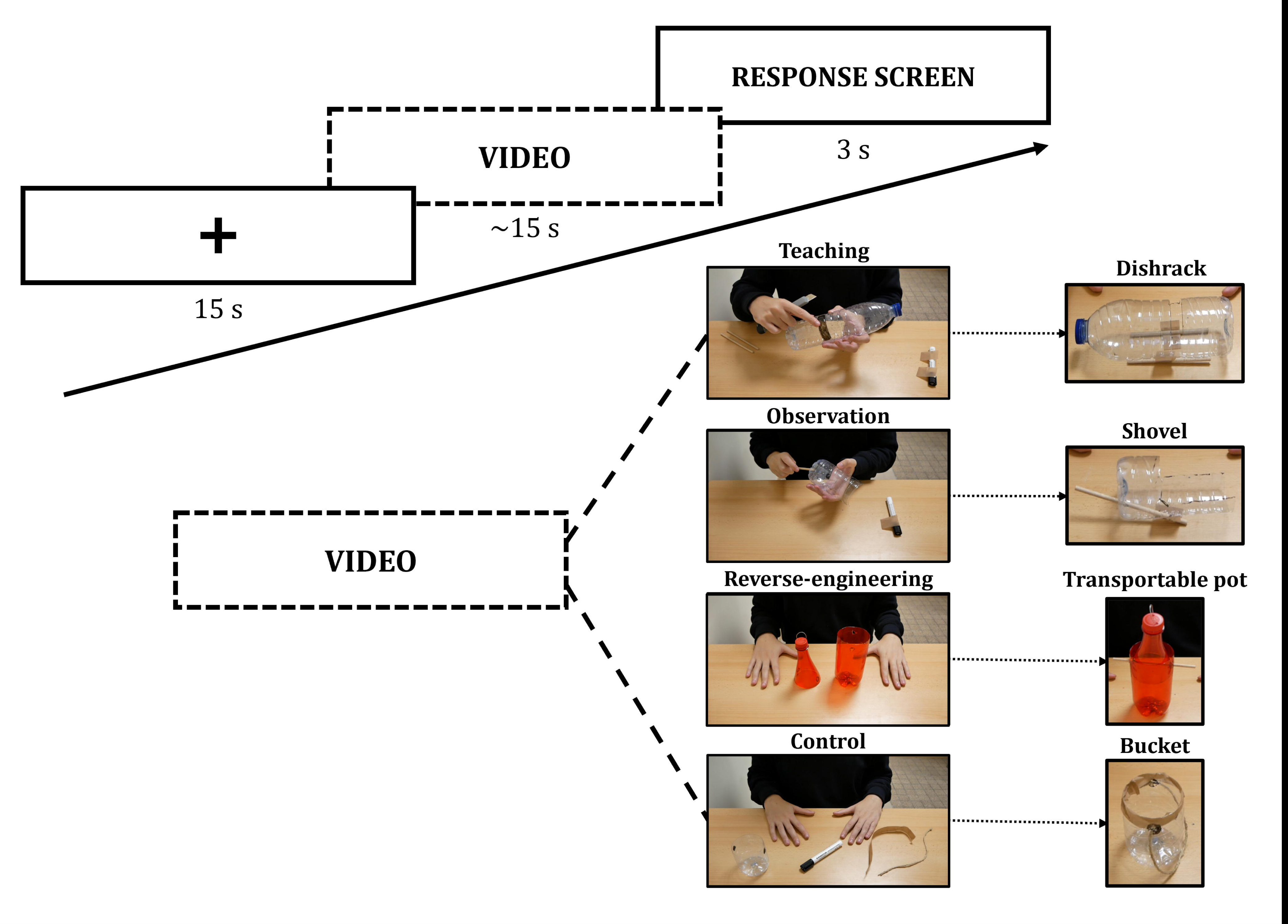

## Materials and methods

### Participants

Thirty healthy participants were enrolled in the study. Inclusion in the final sample required that head motion during scanning did not exceed 0.5mm displacement (i.e., framewise displacement) between consecutive volumes on 90% of volume, resulting in one male participant being excluded on this criterion. Another participant was not scanned due to COVID-related complications. Overall, twenty-height participants (*Mage* = 20.3, *SD* = 2.48; 23 females; all right-handers) completed the study. Sample size was determined based on previous studies that have observed reliable effects of the activations in the technical-reasoning network as well as the tool-use∩action observation network (Lesourd, Afyouni, et al., 2023; Lesourd, Reynaud, et al., 2023; Schubotz et al., 2014). All participants were voluntary and signed written consent. The study was in line with the Declaration of Helsinki and was approved by the French Ethics Committee (N° ID-RCB: 2020-102115-34). Participants received incentives to compensate for their time.

### Stimuli

Three experimental conditions were used in the present study, namely, Reverse engineering, Observation and Teaching. In each condition, participants saw a video representing a scene in which an actor made a tool. **Fig. 1** shows the time course of the experiment along with a screenshot from a video stimulus for each condition. In the Teaching condition, the actor made a tool and communicated non-verbally to explain the process. The same goes for the Observation condition, but without any communication. For the Reverse engineering condition, the actor put forward successively three different making steps of the same tool, starting from the first step onward to the finished tool. We also created a control condition, where the actor just moved around the tools and materials available from the top to the bottom of the table. These tools and materials were matched to the ones used to make the tools. All the videos are available at https://osf.io/6rsqf/.

In total, they were 10 different tools being made across all three conditions, resulting in 30 videos. The scenes were filmed using a Panasonic Lumix G80 camera. The camera was at the back of the table, 50 cm above and tilted at a 30° angle to film from a high-angle spot. Only the arms, hands and torso of the actor were visible, along with the table upon which they were placed. The tools and materials were always placed on the opposite side of the table from the hands of the actor, at the furthest possible distance to still be visible and on the screen. At the beginning of all videos, the hands of the actor were already on the table just in front of him. After some delays that varied between videos, the actor started moving. In the Observation and Teaching conditions, the actor’s hands and arms moved towards the various materials and started the making process. After finishing it, the actor placed the tool on the table so that it was clearly visible to the camera, and then placed his hands in the same way as they were at the beginning of the video. For the Reverse engineering condition, the first step of the making process was already on screen when the video started, and the actor simply moved it aside after 5 seconds while bringing the next step and so forth until the last step was on screen. We directly show the first step to minimize the manipulation of tools and materials in this condition. While the actor was doing nothing, his hands were always in the starting position (i.e., firmly put on the table in front of him).

All videos had a duration of about 15 seconds (*M*_duration_ = 15.15, *SD* = 0.65, *min* = 15, *max* = 17.05; with 59 frames per second) and a resolution of 1280 x 720 pixels. All video editing was performed using the iMovie software (available from https://www.apple.com/fr/imovie/). Editing consisted only of cutting the video so that it started and ended with the actor’s hands on the table. We played with the duration of these start and end positions so that the video was at least 15 seconds long (e.g., when the tool-making process was shorter than 15 seconds, the videos had longer start and end positions). All videos were tested in a separate experiment to make sure participants understood the different conditions. For this experiment, we recruited 5 participants (*M_age_* = 26.2, *SD* = 1.48; 2 females; all right-handers). All participants were beforehand told about the four conditions and were also shown a set of four dummy videos (one per condition) to familiarize themselves with them. Their task was to recognize the condition of the videos. As expected, all participants had a 100% recognition rate.

### fMRI experiment

Each participant was scanned in a single session with: (i) a first functional run, (ii) a T1weighted anatomical scan, and (iii) a second functional run. There were two functional runs because there were two experiments. The order of experiments was counterbalanced between participants. The functional run for the present experiment contained 40 trials, 10 trials per condition, the control condition included. Each trial started with a video (∼15 s) followed by the task screen (3 s) and ended with a rest screen (15 s), consisting of a white fixation cross (15 s) over a black screen. The task screen consisted of a black screen with “MAKING” written on the left and “TRANSPORT” on the right, in the same position as the response buttons. There was no inter-trial interval. A genetic algorithm was used to optimize the experimental design with regards to contrast estimation (Kao et al., 2009) using the toolbox NeuroDesign (https://neurodesign.readthedocs.io/en/latest/index.html).

### Task

Participants were asked to carefully watch each video. After the video, the response screen was shown. They had to indicate if they saw a tool being made or someone moving items around in the videos. To answer, they had a response box with 4 buttons, where the left one was associated with “tool making” and the right one was “item transport”. All three social learning conditions were considered as “tool making”, even Reverse engineering. This task was implemented to maintain the participants’ attention during the viewing of the videos. To ensure the participants understood the instructions during the fMRI sessions, we instructed them before entering the scanner. Because the function of the tool being made in the video was sometimes opaque, we added a familiarization step in the instructions. The instructions consisted of a presentation of the fMRI scanner, the task and instructions. In the familiarization step, the participant saw 10 videos, one per tool, where a demonstrator used the tool in an everyday-life context. They were asked to name the tool or its function and instructed that there was no wrong answer. After the participant was comfortable with the material, the scanning session began.

### fMRI data acquisition

Imaging data were acquired on a 3T Siemens Prisma Scanner (Siemens, Erlangen, Germany) at CERMEP (Lyon, France). A 64-channel head coil was used. Blood-Oxygen Level Dependent (BOLD) images were recorded with T2*-weighted echo-planar images (EPI) acquired with the multi-band sequence. Functional images were all collected as oblique-axial scans aligned with the anterior commissure– posterior commissure (AC–PC) line with the following parameters: 960 volumes per run, 57 slices, TR/TE = 1400 ms / 30 ms, flip angle = 70°, the field of view = 96 x 96 mm^2^, slice thickness = 2.3 mm, voxel size = 2.3 x 2.3 x 2.3 mm^3^, multiband factor = 2. Structural T1-weighted images were collected using an MPRAGE sequence (224 sagittal slices, TR/TE = 3000 / 2.93 ms, inversion time = 1100 ms, flip angle = 8°, 224 x 256 mm FOV, slice thickness = 0.8 mm, voxel size = 0.8 x 0.8 x 0.8 mm^3^).

### Preprocessing of fMRI data

Structural T1-weighted images were segmented into tissue type (GM: grey matter, WM: white matter and CSF: cerebrospinal fluid tissues) using the Computational Anatomy Toolbox (CAT12; http://dbm.neuro.uni-jena.de/cat12/) segmentation tool, to facilitate the normalization step. Functional data were analyzed using SPM12 (Wellcome Department of Cognitive Neurology, http://www.fil.ion.ucl.ac.uk/spm) implemented in MATLAB (Mathworks, Sherborn, MA). Preprocessing for univariate analyses included the following steps: (1) realignment to the mean EPI image with 6-head motion correction parameters and unwarping using the FieldMap toolbox from SPM12; (2) co-registration of the individual functional and anatomical images; (3) normalization towards MNI template; and (4) spatial smoothing of functional images (Gaussian kernel with 5 mm FWHM).

### Group analysis

A General Linear Model was created using design matrices containing one regressor (explanatory variable) for each condition (i.e., Teaching, Observation, Reverse engineering, Control) modelled as a boxcar function (with onsets and durations corresponding to the start of each stimulus of that condition) convolved with the canonical hemodynamic response function (HRF) as well as its temporal and dispersion derivatives. Six regressors of non-interest resulting from 3D head motion estimation (x, y, z translation and three axes of rotation) were added to the design matrix. The model was estimated for each participant, also taking into account the average signal in the run. After model estimation, we computed the three simple contrasts at the first level (i.e., social-learning conditions against control condition) that were transferred to a second-level group analysis (one sample t-tests) to obtain the brain regions activated in the Teaching condition (Teaching > Control), in the Observation condition (Observation > Control), in the Reverse engineering condition (Reverse engineering > Control) and between the Teaching and Observation condition (Teaching > Observation) to isolate the pure teaching component. Afterwards, we entered maps of parameter estimates in a one-way ANOVA, with Social-learning (Teaching, Observation, and Reverse engineering) as a within-subject factor, to identify brain regions where BOLD activity significantly differed across the three conditions. We present results maps with a significance threshold set at *p* < .05 with family-wise error (FWE) correction.

### Seed selection and analysis

A total of 11 spherical seed ROIs (radius = 5mm) were created in the MNI standard space (see **Table 1**). We used literature-defined ROIs to ensure the independence of ROI selection. All our seeds were based on fMRI studies (activation peaks). First, we defined each spherical seed involved in the technical-reasoning network: The left area PF, which has been demonstrated central for technical reasoning (Osiurak & Reynaud, 2020; Reynaud et al., 2019) and the left IFG, which has been repeatedly identified as part of the network but whose role in this network remains unclear (Osiurak, Crétel, et al., 2021). Second, we defined each spherical seed involved in the mentalizing network (Gallagher & Frith, 2003): ACC, which supports the perspective-taking component [(Gallagher & Frith, 2003), localizer in (Molenberghs et al., 2016)], bilateral temporal poles, which are associated with knowledge about persons [(Gallagher & Frith, 2003), localizer in (Molenberghs et al., 2016)], the left STS, which is associated with intention understanding [(Gallagher & Frith, 2003), localizer in (Hodgson et al., 2022)]. And the right MTG [localizer in (Hodgson et al., 2022)], which is known to be involved in communicative gestures (Holler et al., 2015) and communicative intent (Redcay et al., 2016). Finally, the tool-use∩action observation network [all taken from (Reynaud et al., 2019)]: The IPS, which is involved in the motor simulation of hand movements and more precisely grasping preparation (Jeannerod, 1994), the bilateral ITG, which is involved in biological motion perception (Culham et al., 1998; Tootell et al., 1995; Watson et al., 1993) and the left MFG, which might be involved in action-goal perception (Reynaud et al., 2019).

**Table 1.**
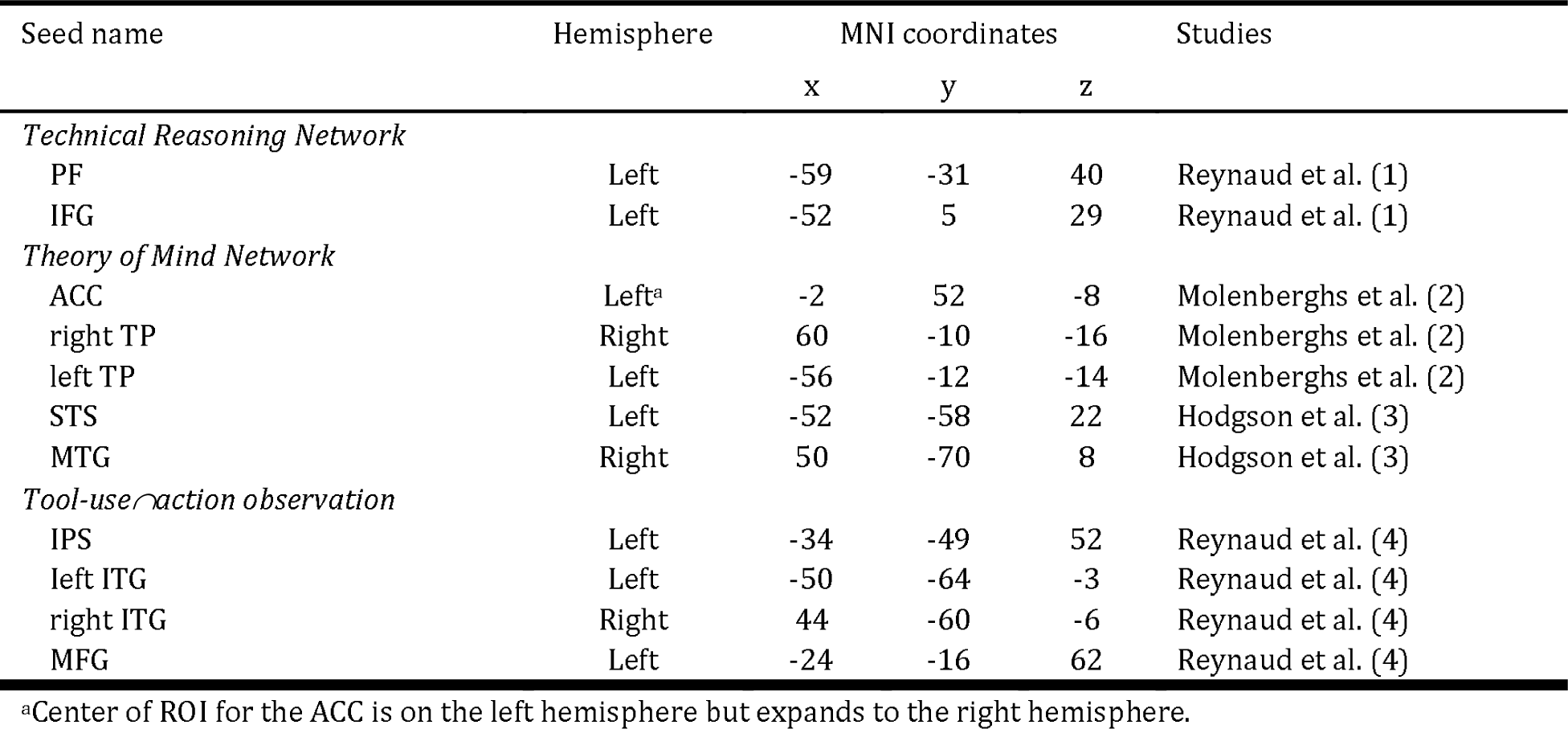
Seed-region locations.

## Results

Participants performed the task correctly as evidenced by the number of correct responses (*M* = 97.59, *SD* = 3.15, *Min* = 87.50, *Max* = 100). For the fMRI results, we first computed the Teaching>Control (**Fig. 2a**), Observation>Control (**Fig. 2b**) and Reverse engineering>Control (**Fig. 2c**) contrasts. The Teaching>Control and the Observation>Control contrasts, but not the Reverse engineering>Control contrast, showed activations in the brain regions associated with the technical-reasoning network (i.e., the left area PF in both contrasts and the left IFG in the Teaching>Control contrast). The brain regions associated with the mentalizing network were recruited neither in the Reverse engineering>Control contrast nor in the Observation>Control contrast, but we found an activation of the right MTG in the Teaching>Control contrast. The involvement of the right MTG in the Teaching>Control contrast was reported in the Teaching>Observation contrast (**Fig. 2d**). We also found activation in the brain areas that support the tool-use∩action observation network in the three experimental conditions (i.e., bilateral ITG and IPS). Finally, we obtained not foreseen activations of the right area PF in both the Teaching>Control contrast and the Observation>Control contrast.

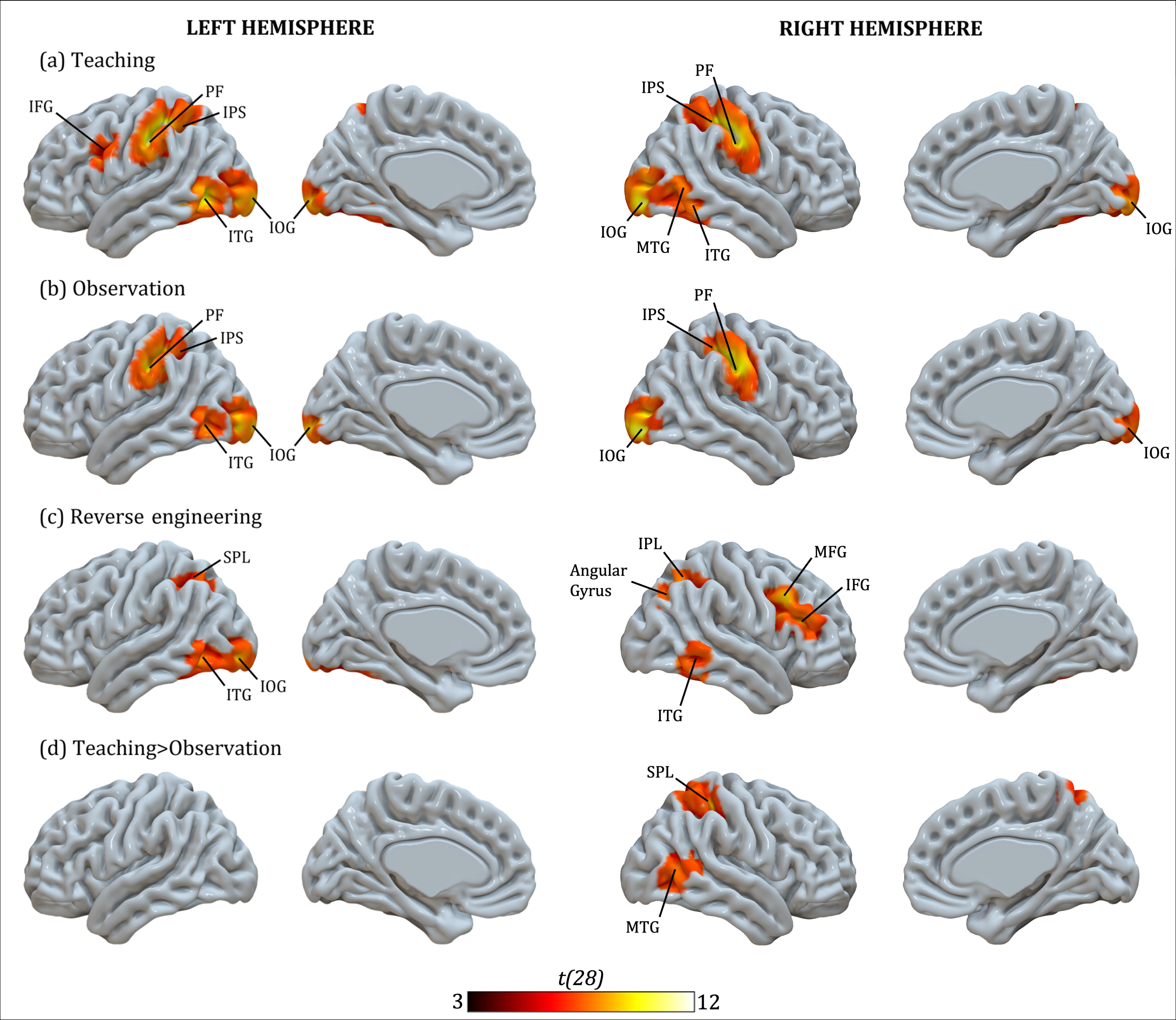

To go deeper into the analysis, we computed a repeated-measures ANOVA based on the individual maps of parameter estimates for the three conditions (Teaching, Observation and Reverse engineering) with Condition as a within-subject factor. The ANOVA showed several regions that differed between the three conditions, namely bilateral occipital cortex, bilateral IPL, left IFG, bilateral SPL, right STG, bilateral MTG, bilateral ITG, left supramarginal gyrus (SMG), and right IFG. Based on these results, we computed posthoc comparisons between conditions in regions of interest (ROI) that showed activation in the ANOVA. As can be seen in **Fig. 3a**, we defined the ROI as aforementioned brain regions associated with the technical-reasoning network (i.e., left area PF and left IFG), the mentalizing network (i.e., ACC, TP, STS, and right MTG) and the tool-use∩action observation network (i.e., IPS, ITG, and left MTG). The list and coordinates of these ROIs are given in **Table 1**. Here we focused on the left area PF, the left IFG, the right MTG, the left IPS, the left ITG and the right ITG as they elicited activation in the ANOVA. Our post hoc tests consisted of *t*-tests on mean activations inside the ROI between the three social-learning conditions (**Fig. 3b-g**). The ROI derived from left PF Teaching (*p* < .001) and Observation (*p* < .05) showed greater activation than Reverse engineering. No difference was observed between Teaching and Observation (**Fig. 3b**). The same pattern was found in the left IFG ROI (both *p* < .001; **Fig. 3c**). For the right MTG ROI (**Fig. 3d**), Teaching elicited greater activation than Observation (*p* < .001) and Reverse engineering (*p* < .001), and Observation greater activation than Reverse engineering (*p* < .001). Teaching preferentially activated the left IPS (**Fig. 3e**) compared to Reverse engineering (*p* < .01). No other statistical difference was reported for the left IPS. Finally, for the left ITG ROI (**Fig. 3f**) and the right ITG ROI (**Fig. 3g**), Teaching showed greater activation than Observation (both *p* < .05) and Reverse engineering (both *p* < .001), and Observation greater activation than Reverse engineering (both *p* < .05). Finally, even if the right area PF was not one of our ROI, we conducted post-hoc *t*-tests to examine the influence of social-learning conditions on its level of activation (**Fig. 3h**). We found that Teaching elicited greater activation than Observation (*p* < .05) and Reverse engineering (*p* < .001), and Observation greater activation than Reverse engineering (*p* < .001).

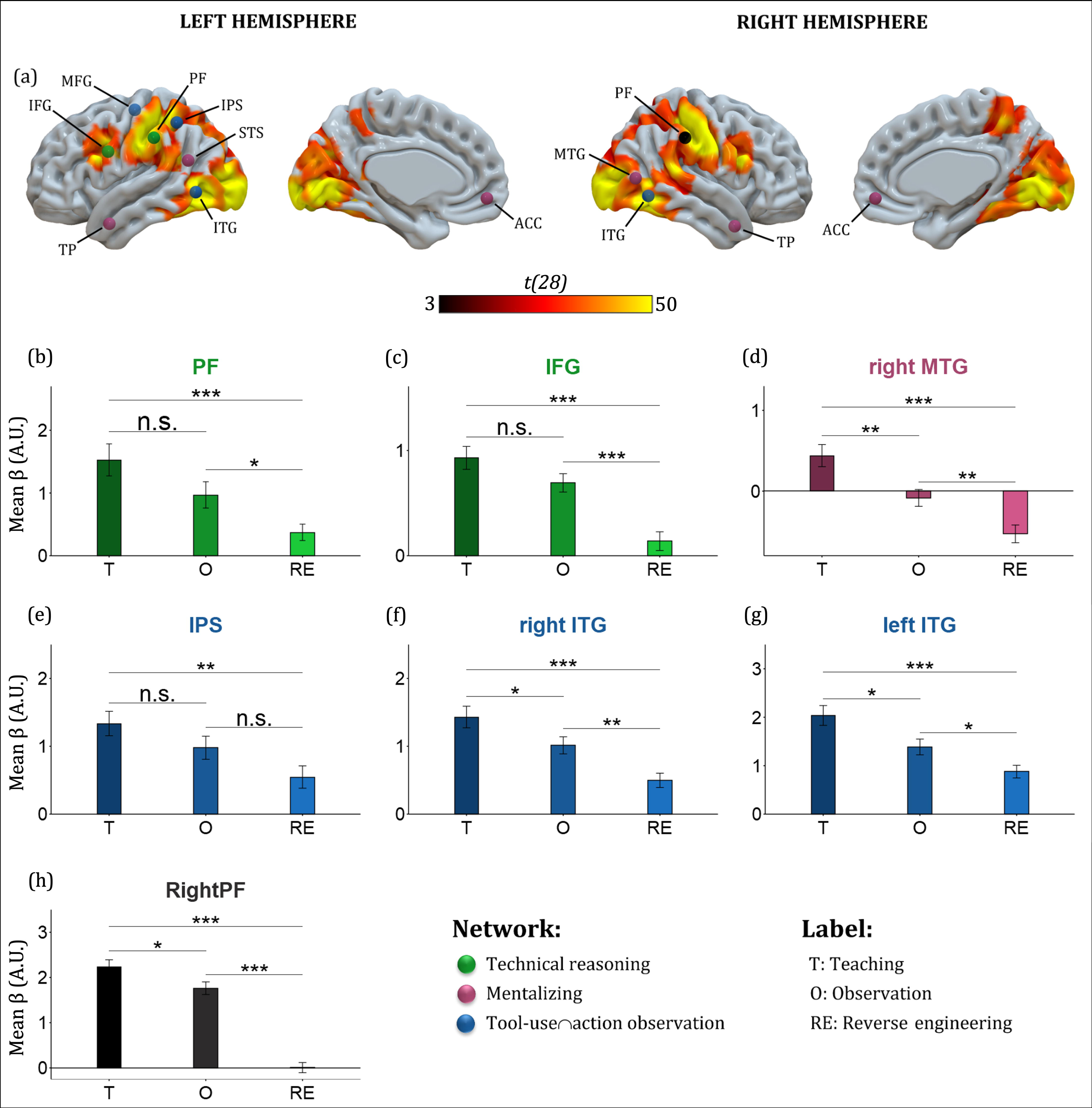

## Discussion

The present study aimed to investigate the neurocognitive origins of social learning during tool-making episodes. To do so, we presented participants with tool-making videos in three social-learning conditions, namely, Reverse engineering, Observation and Teaching. According to the technical-reasoning hypothesis (Bluet et al., 2022; Osiurak, Claidière, & Federico, 2022; Osiurak & Reynaud, 2020), the technical-reasoning network should be recruited in the three social-learning conditions and the mentalizing network only in the Teaching condition. These predictions were partly confirmed by our results. We did find the involvement of the technical-reasoning network in both the Teaching condition and the Observation condition, and we reported the activation of the right MTG, which is part of the mentalizing network, only in the Teaching condition. However, the technical-reasoning network was not activated in the Reverse engineering condition. We will discuss these findings in turn in the next sections.

The first key finding is the recruitment of the technical-reasoning network (i.e., the left area PF and the left IFG) in both the Teaching condition and the Observation condition. Previous findings have indicated that this network is activated when people watch others use tools. We extend these findings by demonstrating that this network is also recruited during the observation of someone making or teaching how to make a tool, thereby drawing a link between technical reasoning, tool making, social learning and, as a result, CTC. As mentioned above, recent accounts have perceived an over-emphasis on the social dimension of the phenomenon [e.g., (Whiten, 2022)], which has led some to minimize the role played by causal understanding/technical reasoning in CTC (Boyd et al., 2011; Derex et al., 2019; Henrich, 2016). Here we show that the ability to reason about our physical world is spontaneously recruited when someone observes a model making or teaching how to make a tool. In these social-learning episodes, technical reasoning allows us to understand the mechanical mechanisms underlying the use or making of tools as well as to infer and predict the goal of each step used to make it. These inferences are essential for the reliable transmission of knowledge related to tool making. In a nutshell, technical reasoning might contribute not only to the innovation component of CTC but also to its high-fidelity transmission component (Osiurak, Claidière, & Federico, 2022).

An unexpected, interesting result is the bilateral activation of the area PF. This bilateral activation is not consistent with the tool-use observation network, in which only the left area PF is involved (Reynaud et al., 2019). It is more consistent with the action observation network, which is distributed bilaterally (Van Overwalle & Baetens, 2009). One possibility is that both hands were visible in our videos, whereas in tool-use videos only the right hand is commonly in action or even visible. Although the right area PF is not known to be a key region of the motor-control system, this remains a viable interpretation. Another possibility is that both tool use and tool making need reasoning about nonspatial, physical object properties (e.g., solidity), which are crucial for generating physical forces (e.g., lever) or mechanical actions (e.g., lever). This aspect might be more left-lateralized and very specific to technical reasoning. By contrast, tool making might need reasoning more about spatial properties than tool use, because it involves assembling objects based on pure spatial dimensions (e.g., orientation). Support for this possibility comes from a recent study that showed that the cortical thickness of the left area PF predicted physical-reasoning performance whereas visuospatial-reasoning performance was predicted by the cortical thickness of both left and right areas PF (Federico et al., 2022). Further studies should try to explore this question.

The second key finding is that isolating the pure teaching component (i.e., Teaching>Observation contrast) led us to find preferential activation of the right MTG, which is part of the mentalizing network. Considering that the right MTG is known for its involvement in the perception of biological motion (Hodgson et al., 2022), this finding is not surprising. However, this region has also been reported to support our understanding of the intention to communicate, also called communicative intent [e.g., (Holler et al., 2015)], which is crucial for teaching. When someone tries to transmit the understanding of a physical phenomenon to another individual, they must first make it clear that that they intend to communicate. In our Teaching condition, this intention was transmitted through non-verbal cues (i.e., pointing and gestures), which helped the model to focus on important elements and steps of the tool-making process. We posit that these cues amplified the learner’s technical reasoning by showing directly where the attention needed to be put. Focusing attention on the crucial parts of the process means spending more time reasoning about them, but it also means not spending time and resources on irrelevant aspects. Thus, the key role of teaching in CTC might be to improve learners’ technical-reasoning skills. In other words, while technical reasoning might be central to CTC, teaching might be part of what makes CTC unique in humans. This specific role of the right MTG is also following the concept of intentional teaching (O’Madagain & Tomasello, 2022), a supposedly unique human form of teaching that allows us to share knowledge more efficiently. This finding also echoes the concept of natural pedagogy (Csibra & Gergely, 2006, 2009), which describes an innate ability for teaching (and being taught), and, therefore, an ancient ability in our evolutionary history. According to the natural pedagogy hypothesis, infants at an early age can capture ostensive signals that indicate that another individual is communicating with them. We propose that this capacity is rooted in the right MTG, which corroborates studies showing that children suffering from autistic disorder both meet difficulties in understanding communicative intent and have abnormal activation of the right MTG [e.g., (Zilbovicius et al., 2006)]. Taken together with our results, these findings suggest an adaptive role of the right MTG for teaching and encourage future research to investigate this link.

It is noteworthy that we did not find any activation of the other brain regions engaged in the mentalizing network (notably ACC and TP) in the Teaching condition. ACC is known to contribute to perspective-taking and TP to the storage of semantic information about persons. Both cognitive components characterize “theory of mind” and might be recruited when people teach instead of when they are taught. This can explain why no activation was found here for these regions in the Teaching condition. Future studies should explore this question by investigating the brain network at work during teaching. We hypothesize that activation of ACC and TP should be found when people teach.

The third finding is that the Reverse engineering condition did not elicit any regions of the technical-reasoning network. This surprising result can be understood in two ways. First, learning via reverse engineering does not require technical reasoning. This is contradictory to several previous studies that have stressed the role of causal understanding/technical reasoning in reverse engineering conditions (Osiurak, Claidière, Bluet, et al., 2022; Zwirner & Thornton, 2015). In broad terms, this interpretation is plausible but unlikely. Second, it could indicate that we failed to create an experimental condition engaging reverse engineering inside the scanner. Reverse engineering is defined as watching a technology and trying to understand how it was made, potentially by manipulating it or by scrutinizing its aspects (Osiurak & Reynaud, 2020). In our experiment, we showed different steps of the making process to the participants for a few seconds at a time, which could be different from a true reverse engineering condition. Moreover, reverse engineering usually implies deconstructing the technology to understand how it was made (e.g., unfolding a paper plane to grasp how it was built), which was not possible for our participants inside the scanner. Furthermore, albeit they knew the tools being built beforehand, the lack of a clear end goal in the video might have made it hard for participants to understand the reason behind the difference in each step shown in the video, and overall make the condition confusing. Finally, the task was to watch the video and respond to whether it depicted a tool-making episode or not. Hence, participants were not forced to understand the making process. Because the setup for the videos in the Control condition were very distinct from the three experimental conditions, participants did not need to pay attention to the whole video to respond. To control for this issue, we ran another analysis where only the first half of the videos was considered. No significant changes were found with this analysis. A solution for studying reverse engineering inside a scanner can be to project a 3D image of a technology and to allow the participant to rotate and zoom on the image. Another solution could be to also investigate reverse engineering using 3D videos of a tool being deconstructed step by step. Otherwise, an alternative would be to change the task, by asking the participant to watch closely the video so that he could make the technology after the scanning session for example. Future research is needed to design a reverse engineering condition suited to neuroimaging conditions.

## Data and code availability

The data that support the findings of the study and the codes used in this paper are available at https://osf.io/6rsqf/.

## Author contributions

A.B., E.R., and F.O. designed the research; A.B., G.F., C.B. F.L., D.I., and Y.R. performed the research. A.B., A.F., and E.R. analyzed data; A.B. wrote the manuscript; E.R., G.F., C.B., M.L., Y.R., and F.O. reviewed and edited the manuscript; F.O., E.R., M.L., and Y.R. acquired the financial support for the project.

## Acknowledgements and Funding Information

This work was supported by grants from the French National Research Agency (ANR; Project TECHNITION: ANR-21-CE28-0023-01; F.O., E.R., M.L., and Y.R.) and the Région Auvergne-Rhône-Alpes (NUMERICOG-2017-900-EA 3082 EMC-R-2075; F.O. and E.R.).

