## Supplementary tables for "The technical-reasoning network is recruited when people observe others make or teach how to make tools: An fMRI study"

**This PDF file includes:**

Tables S1 to S5

| **Table S1.** Local maxima of activation clusters (MNI stereotactic coordinates) for the individual contrasts Teaching>Control, Observation>Control, Reverse engineering>Control and Teaching>Observation | | | | | | | |
| --- | --- | --- | --- | --- | --- | --- | --- |
| Brain region | Hemisphere | Peak MIN | | | Cluster size | T-value | PFWE |
|  |  | x | y | z |  |  |  |
| *Teaching>Control* |  |  |  |  |  |  |  |
| Postcentral Gyrus – PF^a^ | Right | 49 | -18 | 38 | 909 | 16.13 | < .001 |
| Postcentral Gyrus – PF^a^ – IPS^a^ | Left | -34 | -39 | 50 | 821 | 15.71 | < .001 |
| IOG | Right | 30 | -89 | 5 | 505 | 14.84 | < .001 |
| MOG | Left | -29 | -92 | -3 | 477 | 12.83 | < .001 |
| MTG – ITG^a^ | Left | -45 | -62 | -1 | 453 | 11.22 | < .001 |
| ITG – MTG^a^ | Right | 46 | -55 | -12 | 284 | 11.15 | < .001 |
| IFG | Left | -47 | 7 | 27 | 104 | 9.80 | < .001 |
| *Observation>Control* |  |  |  |  |  |  |  |
| Postcentral Gyrus – PF^a^ | Right | 51 | -18 | 38 | 662 | 14.58 | < .001 |
| Postcentral Gyrus – PF^a^ – IPS^a^ | Left | -41 | -30 | 43 | 484 | 11.98 | < .001 |
| IOG | Right | 28 | -89 | -8 | 462 | 15.57 | < .001 |
| IOG | Left | -31 | -89 | -5 | 374 | 12.87 | < .001 |
| IOG – ITG^a^ | Left | -43 | -66 | -3 | 155 | 9.26 | < .001 |
| *Reverse engineering>Control* |  |  |  |  |  |  |  |
| IFG – MFG^a^ | Right | 46 | 16 | 36 | 455 | 11.40 | < .001 |
| ITG | Left | -43 | -53 | -8 | 197 | 12.20 | < .001 |
| Angular Gyrus | Right | 33 | -60 | 48 | 168 | 9.86 | < .001 |
| IPL | Left | -38 | -46 | 41 | 133 | 8.76 | < .001 |
| ITG | Right | 49 | -48 | -15 | 128 | 11.24 | < .001 |
| Cerebellum | Left | -11 | -78 | -35 | 119 | 10.24 | < .001 |
| IOG | Right | -31 | -87 | -8 | 114 | 13.45 | < .001 |
| *Teaching>Observation* |  |  |  |  |  |  |  |
| Postcentral Gyrus | Right | 28 | -34 | 54 | 191 | 11.38 | < .001 |
| MTG | Right | 44 | -60 | 11 | 143 | 9.51 | < .001 |
| All results are thresholded at *p* < .05 (FWE, cluster level).  Brain region labels are given according the aal atlas.  ^a^ Regions that are part of our ROI but they are not the main peak in the cluster they are parts of.  IPS: inferior parietal sulcus; IOG: inferior occipital gyrus; MOG: middle occipital gyrus; MTG: middle temporal gyrus; ITG: inferior temporal gyrus; IFG: inferior frontal gyrus; MFG: middle frontal gyrus; IPL: inferior parietal lobule. | | | | | | | |

| **Table S2.** Region activated in one-way repeated-measure ANOVA | | | | | | | |
| --- | --- | --- | --- | --- | --- | --- | --- |
| Brain region | Hemisphere | Peak MIN | | | Cluster size | F-value | PFWE |
|  |  | x | y | z |  |  |  |
| Lingual Gyrus | Bilateral | 10 | -71 | -5 | 2168 | 315.39 | < .001 |
| IOG – ITG^a^ – MTG^a^ | Right | 28 | -87 | -8 | 2047 | 297.65 | < .001 |
| IOG – ITG^a^ | Left | -31 | -89 | -5 | 2035 | 247.95 | < .001 |
| IPL – PF^a^ – IPS^a^ | Left | -50 | -25 | 38 | 1357 | 200.83 | < .001 |
| IPL – PF^a^ | Right | 53 | -21 | 38 | 1122 | 165.33 | < .001 |
| IFG – MFG^a^ | Left | -50 | 9 | 27 | 374 | 106.77 | < .001 |
| Angular Gyrus | Left | -24 | -76 | 34 | 343 | 78.20 | < .001 |
| IFG | Right | 51 | 11 | 25 | 297 | 114.27 | < .001 |
| STG | Right | 65 | -39 | 15 | 288 | 68.14 | < .001 |
| Postcentral Gyrus | Right | 5 | -48 | 54 | 170 | - | < .001 |
| SPL | Right | 26 | -62 | 34 | 147 | - | < .001 |
| SMG | Left | -59 | -41 | 22 | 146 | - | < .001 |
| All results are thresholded at *p* < .05 (FWE, cluster level).  Brain region labels are given according the aal atlas.  ^a^ Regions that are part of our ROI but they are not the main peak in the cluster they are parts of.  IOG: inferior occipital gyrus; ITG: inferior temporal gyrus; MTG: middle temporal gyrus; IPL: inferior parietal lobule; IPS: inferior parietal sulcus; IFG: inferior frontal gyrus; MFG: middle frontal gyrus; STG: superior temporal gyrus; SPL: superior parietal lobule; SMG: supramarginal gyrus. | | | | | | | |

| **Table S3.** Seed-region locations |  |  | | |  |
| --- | --- | --- | --- | --- | --- |
| Seed name | Hemisphere | MNI coordinates | | | Studies |
|  |  | x | y | z |  |
| *Technical Reasoning Network* |  |  |  |  |  |
| PF | Left | -59 | -31 | 40 | Reynaud et al. (1) |
| IFG | Left | -52 | 5 | 29 | Reynaud et al. (1) |
| *Theory of Mind Network* |  |  |  |  |  |
| ACC | Left^a^ | -2 | 52 | -8 | Molenberghs et al. (2) |
| right TP | Right | 60 | -10 | -16 | Molenberghs et al. (2) |
| left TP | Left | -56 | -12 | -14 | Molenberghs et al. (2) |
| STS | Left | -52 | -58 | 22 | Hodgson et al. (3) |
| MTG | Right | 50 | -70 | 8 | Hodgson et al. (3) |
| *Tool-use∩action observation* |  |  |  |  |  |
| IPS | Left | -34 | -49 | 52 | Reynaud et al. (4) |
| Ieft ITG | Left | -50 | -64 | -3 | Reynaud et al. (4) |
| right ITG | Right | 44 | -60 | -6 | Reynaud et al. (4) |
| MFG | Left | -24 | -16 | 62 | Reynaud et al. (4) |
| ^a^Center of ROI for the ACC is on the left hemisphere but expands to the right hemisphere. | | | | | |

| **Table S4.** Post-hoc *t*-test results | | | |
| --- | --- | --- | --- |
| Seed name | Teaching>Observation | Teaching>Reverse engineering | Observation>Reverse engineering |
| PF | .097^n.s.^ | .000^***^ | .020^*^ |
| IFG | .096^n.s.^ | .000^***^ | .000^***^ |
| IPS | .155^n.s.^ | .002^**^ | .074^n.s.^ |
| left ITG | .016^*^ | .000^***^ | .018^*^ |
| right ITG | .045^*^ | .000^***^ | .003^**^ |
| MTG | .004^**^ | .000^***^ | .005^**^ |
| right PF | .029^*^ | .000^***^ | .000^***^ |
| n.s., not significant.  * *p* < .05  ** *p* < .01  *** *p* < .001 | | | |

| **Table S5.** List of all tools being made in the video stimuli | | |
| --- | --- | --- |
| Name | Description | Photo |
| Bucket | A safe bucket for children to carry toys. | 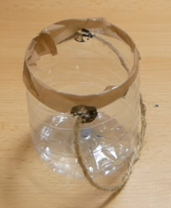 |
| Container | A paper tray that can contain, for example, food. | 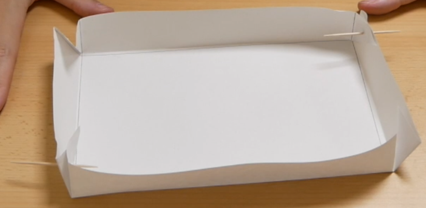 |
| Dishrack | A dishrack for one plate. | 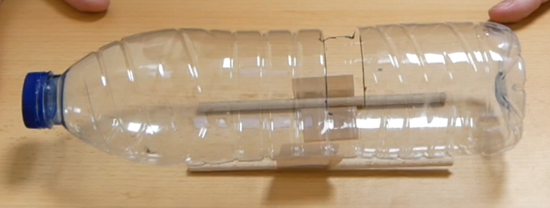 |
| Kazoo | A small musical instrument that produces a buzzing sound. | 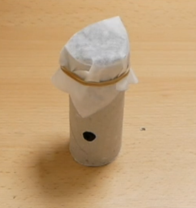 |
| Rake | A small rake for a Zen garden. | 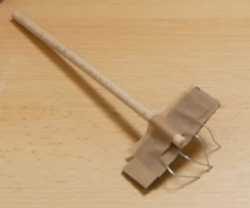 |
| Roll holder | A roll holder for small roll that can be hung. | 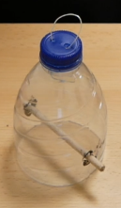 |
| Seed starter | A seed starter with a water tank connected to a pot via a rope, which feeds the plant by capillarity. | 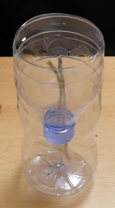 |
| Shovel | A shovel for gardening. | 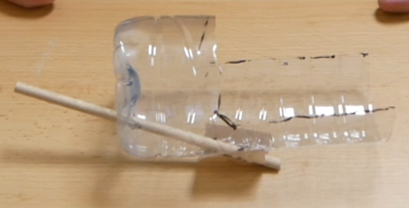 |
| Slingshot | A slingshot that propels objects by pulling on its plastic part. | 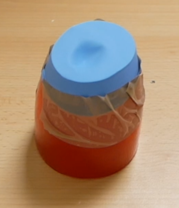 |
| Transportable pot | A pot that can be closed and transported. | 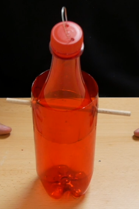 |
